# Immunogenicity and Efficacy of Digitally Immune Optimised H1N1 Vaccine Candidates in Swine and Murine Animal Models

**DOI:** 10.1101/2025.09.12.675937

**Authors:** Joanne Marie M. Del Rosario, Pauline M. van Diemen, Simon Frost, Sneha Vishwanath, Sneha B. Sujit, Sruthika K. Ashokan, Benjamin C. Mollett, Andrew M. Ramsay, Benjamin Simpson, Benedikt Asbach, George W. Carnell, Andrew Chan, Paul Tonks, Nigel J. Temperton, Matthew Davies, John W. McCauley, Rebecca Kinsley, Ralf Wagner, Helen E. Everett, Jonathan L. Heeney

## Abstract

Influenza A virus (IAV) zoonotic transmission and constant evolution in multiple species heightens the risk of emerging novel strains at the human-animal interface. Composite antigens including hemagglutinin (HA), neuraminidase (NA), and matrix-2 (M2) proteins were computationally designed to maximize the breadth of the immune response elicited to human seasonal, pandemic, and zoonotic H1N1 IAVs. Mouse hyperimmune serum raised against these antigens demonstrated broad H1 neutralization and N1 inhibition activity. To enhance immunogenicity, the antigens were combined as a single DNA expression construct (DVX-H1N1). Studies in the well-recognized swine model for human influenza demonstrated that DVX-H1N1 immunization induced broad, neutralizing antibody responses and markedly reduced nasal shedding of viral RNA following challenge with 1A.3.3.2 subclade strain A/swine/England/1353/2009 (H1N1). An effective immune response and reduction in virus shedding was observed in pigs immunized with a whole inactivated virus (WIV) vaccine homologous to the challenge strain but not with a human-origin seasonal WIV vaccine. Overall, we demonstrated broad immunogenicity and efficacy of the DVX-H1N1 vaccine candidate, benchmarked against relevant IAV H1N1 strains *in vitro* and *in vivo* in mice and pigs.

**IMPORTANCE:** The zoonotic potential of swine-origin IAVs is a recognized global health threat. Vaccination remains the most effective intervention against influenza; protecting at the population level by preventing nasal shedding and transmission, but also in individuals by limiting clinical disease, particularly by reducing the severity of lung infection. The World Health Organization (WHO) spearheads biannual surveillance efforts to review evolving virus strains and vaccine antigens at Vaccine Candidate Meetings (VCM) to recommend strain updates for the human seasonal influenza vaccine and for pandemic preparedness purposes. However, the strain selection approach is complex and efficaciousness of seasonal influenza vaccines still varies significantly based on the accurate matching of the predicted strains in circulation with the manufactured vaccine antigens. This emphasizes the need for next-generation influenza vaccines that improve the breadth and longevity of immunity. We describe a computationally optimized DNA vaccine with broad immunogenicity and robust efficacy in the pig model.

## INTRODUCTION

Influenza A virus (IAV) is a zoonotic virus with human epidemic and pandemic potential (1). IAVs are classified according to their envelope glycoproteins, hemagglutinin (HA) and neuraminidase (NA). Only three HA (H1, H2, H3), and two NA (N1, N2) subtypes have resulted in past human pandemics and continuing endemic human infections (2–4).

IAV infection continues to be a global public health burden with the WHO estimating that a billion people experience seasonal influenza annually, with 3-5 million people affected with severe illness resulting in 290,000-650,000 respiratory deaths (5). Vaccination remains the most effective intervention to mitigate the impact of disease. Conventional influenza vaccines can limit clinical disease following infection (6,7) but may not entirely prevent nasal shedding of virus by all vaccinated individuals. Additionally, even when vaccines are antigenically matched to circulating influenza strains, in the US, influenza vaccines conferred on average 40% vaccine effectiveness in the last 15 years (8). Despite immunization, many develop severe influenza that requires medical intervention and in years when vaccine antigens are poorly matched to circulating viruses, efficacy can diminish to as low as 10% (9).

Influenza viruses undergo constant genetic change as low fidelity RNA replication, and the eight-segmented genome allow for the gradual accumulation of genetic mutations (drift) and entire gene segment exchange (shift). Antigenic drift as well as shift events mainly centered on HA and NA lead to antigenic change associated with immune escape and widespread infection as well as transmission between species (10,11) and rare co-infections can result in genetic reassortment with the potential for emergence of viruses with novel gene constellations and biological properties that drive pandemic emergence. Moreover, HA and NA of an influenza subtype can evolve independently of each other. These two Influenza A surface glycoproteins exert opposite functions, with HA facilitating viral attachment, and NA, viral release. **Fig. 1A** shows time trees of HA (left) and NA (right) from the H1N1 pandemic lineage since 2009. The leaves of the time trees of HA and NA from same viral strain are connected by lines. The crossing of the lines suggests different evolutionary trajectories of HA and NA (**Fig. 1A**).

**FIG 1.**
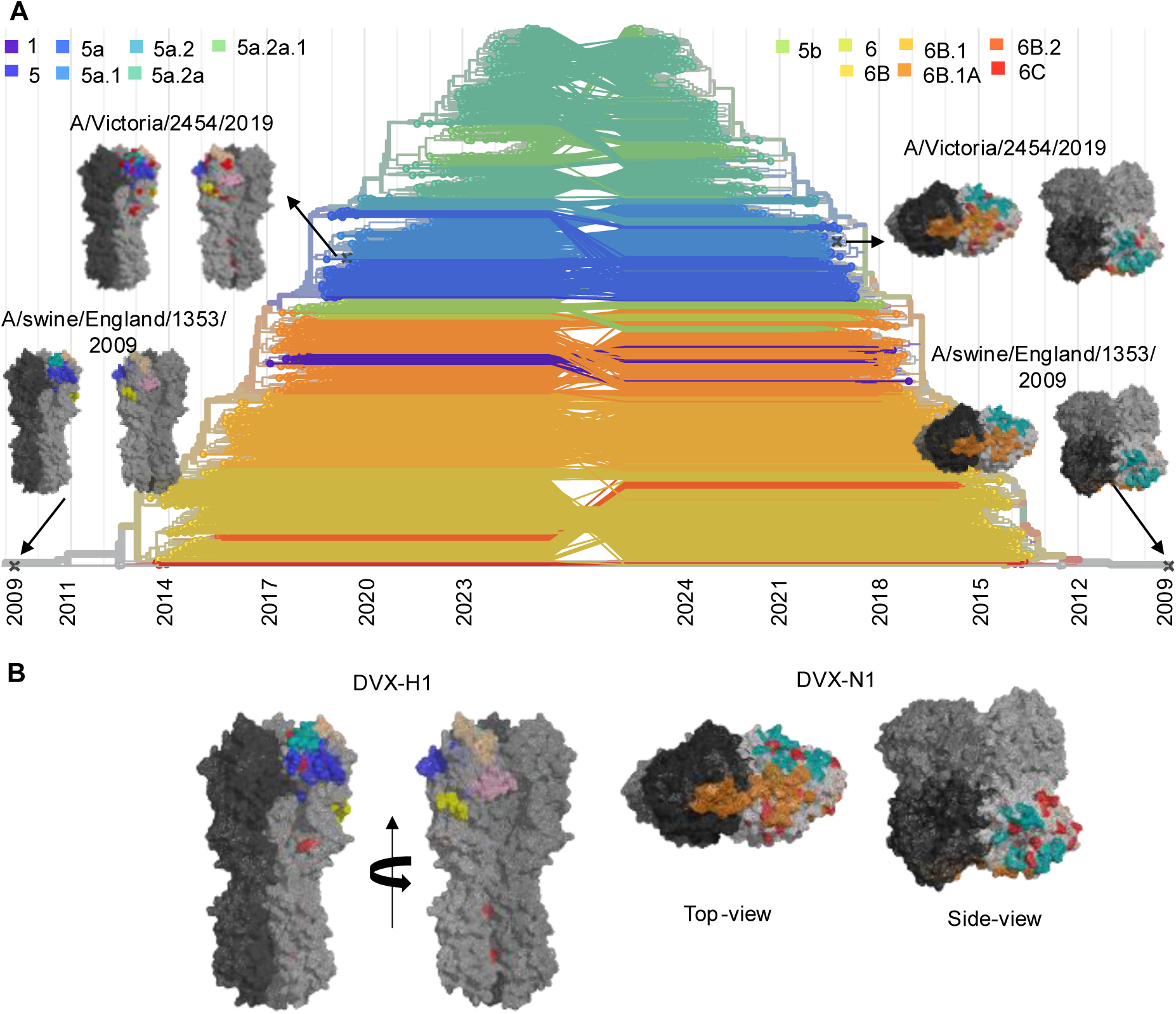
Evolution of Hemagglutanin (HA) and Neuraminidase (NA) of the influenza 1918 H1N1 pandemic lineage. **(A)** A time tree representation of HA (left) and NA (right) from H1N1 pandemic lineage in humans since 2009. All the sequences are represented as the filled circles and colored according to the lineage. A/swine/England/1353/2009 and A/Victoria/2454/2019 are represented as ‘X’ in the time tree to show the evolutionary distance between the two viruses. Structures of the ectodomain of HA and the head domain of NA are shown as surface representations. The residues different between the A/swine/England/1353/2009 and the A/Victoria/2454/2019 are highlighted in red for both HA and NA. **(B)** Structural model of the ectodomain of DVX-H1 and the head domain of DVX-N1. The amino acid differences between DVX-H1/N1 and A/swine/England/1353/2009 are highlighted in red. For both the panels, the antigenic sites – SA, SB, CB, CA1, and CA2 for HA are highlighted as light orange, teal, yellow, pink, and blue, respectively. The receptor binding site for HA is shown as spheres. The antigenic sites of NA are highlighted as orange and teal. The time trees were generated using Nextstrain (70,71). Surface representation of HA (72) and NA (73) were generated and rendered using Pymol Open source v2.5.0 (LLC Schrodinger and Warren DeLano).

IAV host tropism is largely determined by the affinity of the viral HA receptor binding domain (RBD) for sialic acid receptors on the surface of host cells that are required for viral attachment and entry (12). Immune responses directed against the HA, particularly when evaluated using the Hemagglutination Inhibition (HI) test, are used as a surrogate for vaccine-mediated protective efficacy. Additionally, antibodies specific for HA or NA are also known to protect experimental animals from IAV pathogenesis (13–15).

Pigs can serve as important intermediate host species (16) as they are susceptible to infection by IAVs originating from multiple species, largely due to the presence of both α-2,3-and α-2,6-linked sialic acid receptors used by avian and mammalian-origin HA proteins, respectively, for airway cell attachment and infection (7,12,17). Cross-species transmission into pigs, followed by onward transmission and adaptation leads to the establishment of enzootic swine IAV (IAV-S) strains in pig populations. The zoonotic threat posed by IAV-S was exemplified in 2009, when an H1N1 swine virus infected humans and became pandemic (17,18). This H1N1 strain, now designated Clade 1A.3.3.2 in pigs, and H1N1pdm09 in humans, had acquired genetic elements from ancestral swine-origin H1N1 and triple-reassortant lineages of North American origin combined with NA and matrix (M) gene segments originating from a Eurasian avian-like (HA clade 1C) swine virus, which was originally of avian origin (19,20). This new 1A.3.3.2 clade rapidly transmitted within human and pig populations globally and increased genetic diversity, reassortment, and fitness of IAV in both host species. Sporadic human infection by so-called variant (v) swine-origin IAV strains from multiple clades continues to be reported (21). These viruses are enzootic in pig populations in most regions of the world with the underlying potential to cause a human pandemic when allowed to evolve in pigs (22).

The evolving nature of viruses in animals with spillovers to humans highlights the importance of a “One Health” approach to infectious disease control across both human and veterinary medicine to better tackle infectious disease threats, particularly influenza. Benefits include a more effective global surveillance, co-operation and response for the early intervention and prevention of disease emergencies and outbreaks while strengthening pandemic preparedness at the international level (23,24). Lessons learned from the COVID-19 pandemic, require a stronger focus on early detection of zoonotic threats and implementation of intervention strategies (25).

Evaluation of the best targets for vaccination and/or drug development is paramount in determining deployment approaches to tackle influenza prevention and management. Following influenza infection or vaccination, the host response is primarily directed against the HA and NA glycoproteins of the virion envelope (9) with other viral components including the nucleoprotein (NP), non-structural (NS), and matrix proteins (M1 and M2), having less well-characterized roles in the host immune response to infection and protection (12). In the last few years, different vaccine approaches using chimeric HA (26), virus-like particles (VLPs) (15,27,28), consensus based influenza antigens (COBRA) (29), nanoparticles (27,30–32), and nucleic acid vaccines (33,34) have been developed. However, these technologies still require comprehensive and costly global surveillance to provide precise epidemiological information (35,36). Moreover, swine influenza vaccines are more constrained; vaccine antigen updates are limited by regulatory hurdles and given the notable global diversity of IAV-S antigens, may show limited efficacy and elicit low cross-reactive humoral immunity against the heterologous strains that circulate in pig herds (30,37–41).

To address this problem, we undertook a Digitally Immune Optimized Synthetic Vaccine (DIOSynVax) approach that utilizes available sequence data from IAV circulating in animal and human populations. We focused on HA, and NA from H1N1 influenza to maximize coverage of human seasonal, pandemic, and zoonotic IAV strains. Here we demonstrate that antigens generated using DIOSynVax technology (42), delivered by DNA immunization in mice and pigs, produced robust immune responses. A representative vaccine, DVX-H1N1, was highly efficacious when assessed in pigs as a well-established large animal model for human influenza (43) following challenge with a biologically relevant swine Clade 1A3.3.2 challenge virus, A/swine/England/1353/2009 (H1N1) (44).

## RESULTS

### *In-silico* antigen designs

To develop a vaccine capable of eliciting broader immune response across all the lineages of H1N1 influenza, we computationally designed three antigens: an antigen representing hemagglutinin (HA) of H1N1 pandemic lineage - DVX-H1, neuraminidase (NA) of N1 subtype - DVX-N1, and matrix protein 2 (M2) of Influenza A viruses - DVX-M2. Here, only DVX-H1 and DVX-N1 are discussed, and DVX-M2 is discussed in detail elsewhere (45). For designing these antigens, a dataset of non-redundant HA sequences from H1N1 pandemic lineage, and a dataset of non-redundant NA from N1 subtypes, were compiled using NCBI database (46). Sequences reported up to 2018 were used for all the antigens. Due to the evolutionary relatedness between the input sequences and the evolution model used in our pipeline to generate these three antigens, the designs capture both the conserved as well as distinct epitopes of the input sequences (**Fig. 1B**) and are thus expected to generate broad immune responses to known H1N1 viruses.

### DVX antigens elicit broad neutralising responses in the mouse model

The immunogenicity of DVX-H1 and DVX-N1 vaccine candidates expressing individual H1, and N1 antigens (**Table 1**) were evaluated relative to the “empty” vector plasmid control in the mouse model. DNA expressing proteins from the recommended WHO candidate vaccine viruses at the time of DVX antigen design (2018), indicated as H1_ctrl_, and N1_ctrl_, were also included as positive control immunogens.

**Table 1.**
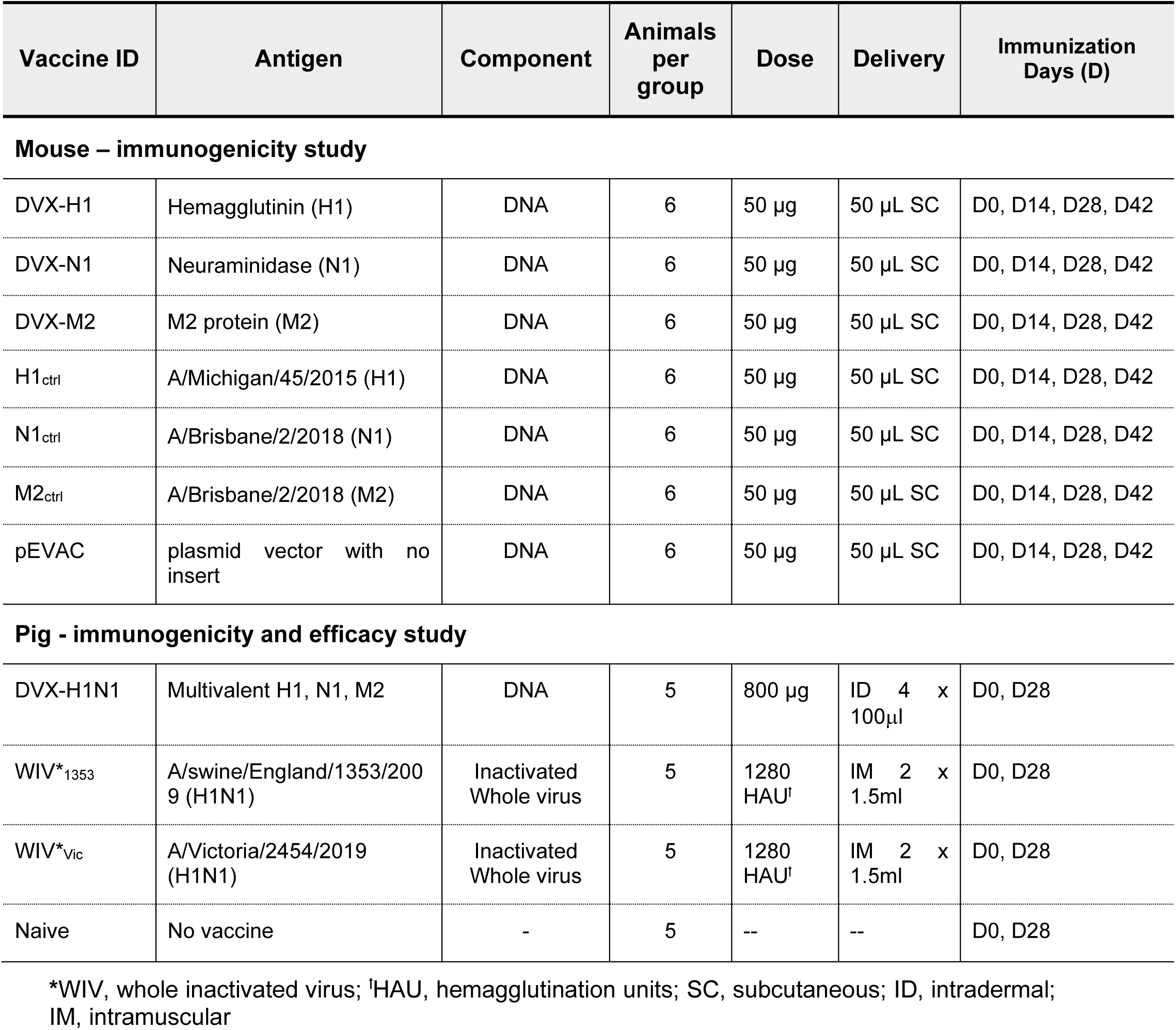
Antigens used in animal studies.

| Vaccine ID | Antigen | Component | Animals per group | Dose | Delivery | Immunization Days (D) |
| --- | --- | --- | --- | --- | --- | --- |
| **Mouse – immunogenicity study** |  |  |  |  |  |  |
| DVX-H1 | Hemagglutinin (H1) | DNA | 6 | 50 µg | 50 µL SC | D0, D14, D28, D42 |
| DVX-N1 | Neuraminidase (N1) | DNA | 6 | 50 µg | 50 µL SC | D0, D14, D28, D42 |
| DVX-M2 | M2 protein (M2) | DNA | 6 | 50 µg | 50 µL SC | D0, D14, D28, D42 |
| H1<sub>ctrl</sub> | A/Michigan/45/2015 (H1) | DNA | 6 | 50 µg | 50 µL SC | D0, D14, D28, D42 |
| N1<sub>ctrl</sub> | A/Brisbane/2/2018 (N1) | DNA | 6 | 50 µg | 50 µL SC | D0, D14, D28, D42 |
| M2<sub>ctrl</sub> | A/Brisbane/2/2018 (M2) | DNA | 6 | 50 µg | 50 µL SC | D0, D14, D28, D42 |
| pEVAC | plasmid vector with no insert | DNA | 6 | 50 µg | 50 µL SC | D0, D14, D28, D42 |
| **Pig - immunogenicity and efficacy study** |  |  |  |  |  |  |
| DVX-H1N1 | Multivalent H1, N1, M2 | DNA | 5 | 800 µg | ID 4 x 100µl | D0, D28 |
| WIV*<sub>1353</sub> | A/swine/England/1353/2009 (H1N1) | Inactivated Whole virus | 5 | 1280 HAU<sup>†</sup> | IM 2 x 1.5ml | D0, D28 |
| WIV*<sub>Vic</sub> | A/Victoria/2454/2019 (H1N1) | Inactivated Whole virus | 5 | 1280 HAU<sup>†</sup> | IM 2 x 1.5ml | D0, D28 |
| Naive | No vaccine | - | 5 | -- | -- | D0, D28 |

\*WIV, whole inactivated virus; <sup>†</sup>HAU, hemagglutination units; SC, subcutaneous; ID, intradermal;
IM, intramuscular

The humoral immune response elicited at the study end after 70 days after four immunizations was evaluated using HA neutralisation pseudotype virus (PV) (**Table 2**) assays. All mice immunized with DVX-H1 developed an immune response against all H1 PV tested (**Fig. 2A**). Neutralizing (IC_50_) titres in mice immunized with the H1_ctrl_ antigen were detected against the majority of the seasonal H1 strains; however, these were absent against H1 PVs representing strains BR/59/07 (HA Clade 1B1.2), sw/GX/13 and sw/HN/18 (HA Clade 1C.2.3) as well as from SC/1/1918, the ancestral 1A virus A/South Carolina/1/1918. The vector control vaccination did not elicit any measurable immune responses, as expected. Additionally, a Hemagglutination Inhibition assay (HI) was used to evaluate HA head domain-specific antibodies against representative seasonal H1N1 strains and variant swine-origin IAV (A(H1)v) viruses that have caused human infections (**Table 3**). Serological responses to H1N1 viruses CA/07/09, MI/45/15, and BR/2/18, all of which are recent human seasonal vaccine antigens, were detected for both DVX-H1 and H1_ctrl_; however, titres were lower following DVX-H1 immunization compared to the homologous viral antigen H1_ctrl_ representing MI/45/15 (**Fig. 2B**). Nonetheless, in terms of breadth, the DVX-H1 antigen elicited a detectable immune response against the A(H1)v viruses Hun/42443/15 and H-H/1572/19 with HA from clade H1C.2.3, while there was no response detected in H1_ctrl_ immunised mice against the A(H1)v viruses. Neither antigen elicited a serological response to the HESS/47/20 (clade 1C.2.2) strain. These data demonstrate the limited strain specificity of a seasonal influenza HA antigen compared to the broad specificity of DVX-H1, as shown by serum neutralization titres using a diverse H1 PV panel compared to H1_ctrl_ DNA (**Fig. 2A**) as well as HI titres against both human-and swine-origin H1N1 viruses from different clades (**Fig. 2B**).

**FIG 2.**
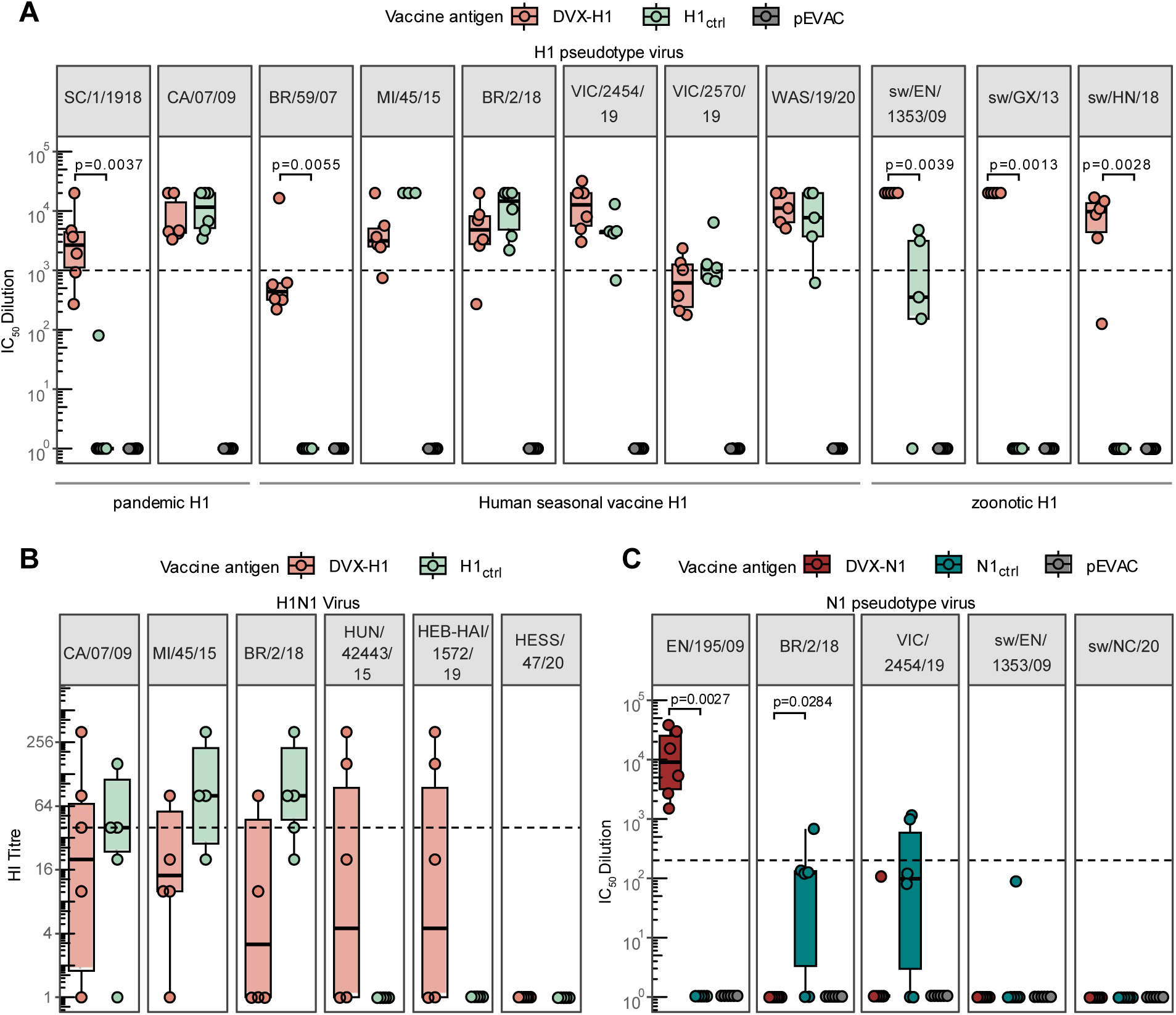
Serological responses of mice immunised with DVX-H1, DVX-N1, H1_ctrl_, N1_ctrl_, and vector control (pEVAC) plasmids. **(A)** Neutralization of H1 pseudotype viruses (PV). Serum samples obtained at 70dpi were tested against a panel of pandemic, seasonal, and zoonotic H1 PV (**Table 2**). Serum neutralizing titres are represented as IC_50_ values (fold dilution of sera that resulted in 50% neutralization of virus). The dashed line indicates an IC_50_ value of 1000, the threshold for a predicted protective immune response in this assay (61,73–76). **(B)** Hemagglutination inhibition (HI) titres against a panel of H1N1 virus strains (**Table 3**). The dotted line indicates an HI titre of 40 which is considered the lower seropositive threshold. **(C)** Inhibition of N1 pseudotype viruses. IC_50_ dilution values were determined via pELLA. The dashed line indicates an IC_50_ value of 200, the predicted baseline for an effective neuraminidase inhibition response in this assay (62). For all plots, the median and interquartile range of individual mouse serum samples are shown (n=6) per vaccination group. Only significant *p* values *(<0.05*) of DVX-H1 vs H1_ctrl_ (**A,B**), and DVX-N1 vs N1_ctrl_ (**C**) as determined by Wilcoxon signed-rank test are shown for clarity. All other p values are tabulated in **Supplementary Table 1**.

**Table 2.** Virus strains and sequences used to generate HA and NA pseudotype viruses.

| Virus Strain | Designation | Abbreviation | Accession number | HA Clade |
| --- | --- | --- | --- | --- |
| **HA** |  |  |  |  |
| A/South Carolina/1/1918 (H1) | pandemic | SC/1/1918 | AF117241 | 1A.1 |
| A/California/07/2009 (H1) | pandemic | CA/07/09 | MN596845 | 1A.3.3.2 |
| A/England/195/2009 (H1) | pandemic | EN/195/09 | GQ166661.1 | 1A.3.3.2 |
| A/Brisbane/59/2007 (H1) | former seasonal | BR/59/07 | CY058487 | 1B.2 |
| A/Michigan/45/2015 (H1) | seasonal pdm09 | MI/45/15 | EPI662594 | 1A.3.3.2 |
| A/Brisbane/2/2018 (H1) | seasonal pdm09 | BR/2/18 | EPI1383389 | 1A.3.3.2 |
| A/Victoria/2454/2019 (H1) | seasonal pdm09I | VIC/2454/19 | EPI1701884 | 1A.3.3.2 |
| A/Victoria/2570/2019 (H1) | seasonal pdm09 | VIC/2570/19 | EPI1741926 | 1A.3.3.2 |
| A/Washington/19/2020 (H1) | seasonal pdm09 | WAS/19/20 | MT331513 | 1A.3.3.2 |
| A/swine/England/1353/2009 (H1) | pandemic, zoonotic | sw/EN/1353/09 | KR701097.1 | 1A.3.3.2 |
| A/swine/Guangxi/BB1/2013 (H1) | zoonotic | sw/GX/13 | KJ725056 | 1C.2.3 |
| A/swine/Henan/SN/10/2018 (H1) | zoonotic | sw/HN/18 | MN416619 | 1C.2.3 |
| **NA** |  |  |  |  |
| A/England/195/2009 (N1) | pandemic | EN/195/09 | GQ166659 | 1A.3.3.2 |
| A/Brisbane/2/2018 (N1) | seasonal | BR/2/18 | EPI1312565 | 1A.3.3.2 |
| A/Victoria/2454/2019 (N1) | seasonal | VIC/2454/19 | EPI1701883 | 1A.3.3.2 |
| A/swine/England/1353/2009 (N1) | pandemic, zoonotic | sw/EN/1353/09 | EPI640886 | 1A.3.3.2 |
| A/swine/North Carolina /A02478985/2020 (N1) | zoonotic | sw/NC/20 | MT020063 |  |

**Table 3.** Panels of virus strains used in the hemagglutination inhibition (HI) assay.

| Virus Strain | Abbreviation | H1N1 Clade information<br>(swine nomenclature extended to human isolates) |
| --- | --- | --- |
| **Mouse panel** |  |  |
| A/California/07/2009 (H1N1)* | CA/07/09 | H1A.3.3.2 human seasonal vaccine strain 2010- 2016 |
| A/Michigan/45/2015 (H1N1)~ | MI/45/15 | 1A.3.3.2 human seasonal vaccine strain 2017- 2019 |
| A/Brisbane/2/2018 (H1N1)~ | BR/2/18 | 1A.3.3.2 human seasonal vaccine strain 2019-2020 |
| A/Hunan/42443/2015 (A(H1)v) | HUN/42443/15 | H1C.2.3 Eurasian avian-like variant strain |
| A/Hebei-Haigang/1572/2019 (A(H1)v) | H-H/1572/219 | H1C.2.3 Eurasian avian-like H1N1 variant strain |
| A/Hessen/47/2020 (H1N1) | HESS/47/20 | 1C.2.2 variant H1N1 strain |
| **Pig panel** |  |  |
| A/Victoria/2454/2019 (H1N1)~ | VIC/2454/19 | H1A.3.3.2 human seasonal vaccine strain 2020-2021, homologous to WIV<sub>VIC</sub> |
| A/England/195/2009 (H1N1)* | EN/195/09 | 1A.3.3.2 variant and human seasonal vaccine candidate |
| A/swine/England/1353/2009 (H1N1)*^ | sw/EN/1353/09 | H1A.3.3.2, challenge strain homologous to WIV<sub>1353</sub> |
| A/swine/England/13C4WA/2022 (H1N1)^ | sw/EN/13C4WA/22 | H1A.3.3.2 Contemporary swine-origin H1N1 strain |

\*Pandemic IAV, ~Human Seasonal Vaccine IAV, ^Zoonotic IAV

Neuraminidase inhibition (NAI) titres elicited at 70dpi following immunization with the DNA construct DXV-N1 were evaluated by pELLA employing N1 PV (**Table 2**). NAI activity was detected against the N1 PV representing the EN/195/09 but not with the other N1 PV tested (**Fig. 2C**). Sera from mice in the N1_ctrl_-immunized group showed some NAI activity against the homologous human seasonal strains BR/2/18 (N1), and against VIC/2454/19 (N1) PV but below the threshold to produce an IC_50_ response. Sera from all groups of mice immunized with DVX-M2 or M2_ctrl,_ did not show significant specific binding activity against M2 peptides as tested via a Luminex Assay as shown previously (45).

### DVX-H1N1 elicits immune responses comparable to whole inactivated virus (WIV) vaccines in pigs

The well-recognized pig model of influenza was used to investigate immunogenicity and efficacy of DVX-H1, DVX-N1, and DVX-M2 combined into a single DNA plasmid, DVX-H1N1, in comparison to whole inactivated virus (WIV) vaccines incorporating the antigens A/swine/England/1353/2009 (WIV_1353_) or A/Victoria/2454/2019 (WIV_Vic_) (**Table 1**). HA and NA of WIV_1353,_ and WIV_Vic,_ have differences in the known antigenic site of HA and NA (**Fig. 1A**). Three groups of pigs (n=5) were immunized by needle-free intradermal administration (Pharmajet) of DVX-H1N1 DNA or intramuscularly with adjuvanted WIV vaccines according to the study schedule indicated (**Fig. 3A**). A fourth group of naive control animals was included. No adverse clinical signs were observed upon immunization.

**FIG 3.**
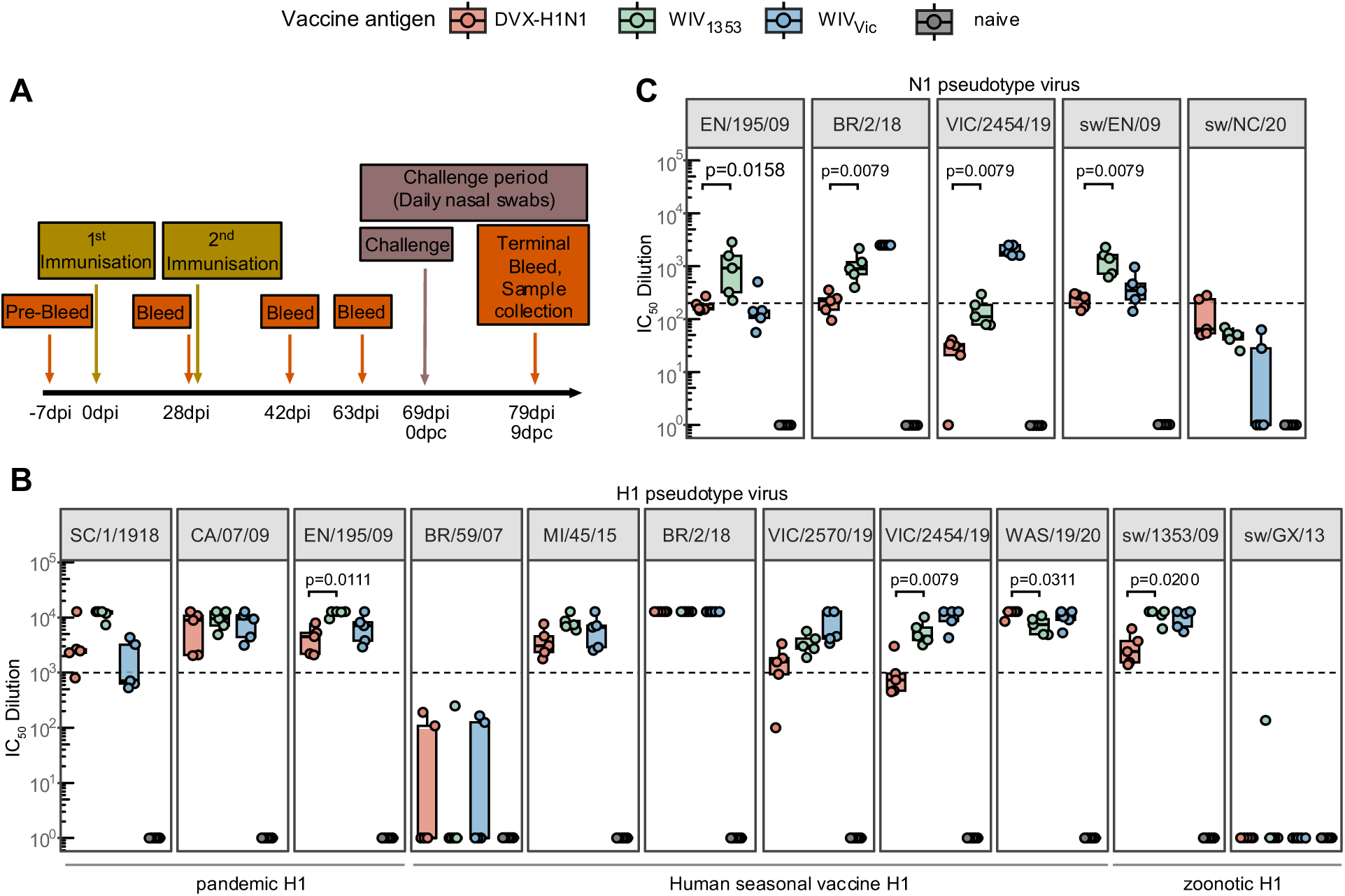
Serological results showing breadth of immune responses of pigs immunized with DVX-H1N1, WIV_1353_, WIV_VIC_, and non-vaccinated group on 42dpi**. (A)** Immunization and sampling schedule of pigs immunized with DVX antigens and control whole inactivated virus (WIV) vaccines prepared from antigens A/swine/England/1353/2009 (WIV_1353_) and A/Victoria/2454/2019 (WIV_Vic_). Serum samples obtained at 42 days post-immunization (dpi) were analzsed. **(B)** Neutralization of H1 pseudotype viruses (PV). Post-immunization sera were tested against a panel of pandemic, seasonal, and zoonotic H1 PV strains (**Table 2**) and neutralizing titres are represented as IC_50_ dilution values. Dashed line indicates an IC_50_ value of 1000. **(C)** Inhibition of N1 pseudotype viruses (**Table 2**). IC_50_ dilution values were determined via pELLA. Dashed line indicates an IC_50_ value of 200. For all plots, the median and interquartile range of individual pig serum samples are shown (n=5) per vaccination group. Only significant *p* values *(<0.05*) of DVX-H1N1 vs WIV_1353_ as determined by Wilcoxon signed-rank test are shown for brevity. All other p values are tabulated in **Supplementary Table 2**.

Serum samples obtained at 42dpi were used to assess the breadth of the humoral responses induced by DVX-H1N1 and WIV immunizations by testing sera against H1 and N1 PV panels (**Table 2**). Broad anti-H1 neutralizing responses (IC_50_≥10^3^) were detected in all vaccinated pigs against the majority H1 PV tested (**Fig. 3B**). There was no neutralizing response detected to sw/GX/13 H1 PV in serum from any group and only a few animals from the DVX-H1N1 and WIV_VIC_ immunized groups showed a serological response to the pre-2009 human seasonal influenza virus BR/59/07 H1 PV, but with titres IC_50_≤10^3^ (**Fig. 3B**). All immunized groups showed IC_50_ NAI activity against all N1 PV tested (**Fig. 3C**). Only titres measured using the sw/EN/1353/09 N1 PV, which is homologous to the WIV_1353_ vaccine, reached an IC_50_≥10^2^. As expected, serum from the WIV_VIC_ immunized group demonstrated significantly higher N1 NAI activity to the homologous human seasonal N1 PV and the N1 PV representing a 2009 vaccine strain of the same lineage, EN/195/09, compared to DVX-H1N1 and WIV_1353_-immunized animals.

### DVX-H1N1 protects pigs from challenge employing both humoral and cell-mediated immune responses

We then investigated the corresponding efficacy afforded by these immunizations (**Fig. 3A**). All pigs were challenged intranasally with the A/swine/England/1353/2009 (H1N1) virus on 69dpi (corresponding to 0 days post challenge (dpc)). Clinical signs were mild or non-existent after inoculation and clinical scores transiently reached no more than 3 out of a possible maximum of 21 (on a daily basis) using an established clinical scoring system. Rectal temperatures showed a slight transient increase in all pigs on 4-5dpc (**Sup. Fig. 1A**) while all animals remained clinically healthy.

Nasal shedding of viral RNA was monitored daily by RT-qPCR. All inoculated animals shed viral RNA, reaching a peak between 2-5dpc (**Fig. 4A**). Both the DVX-H1N1 and homologous WIV_1353_ immunized animal groups showed a markedly lower level of viral RNA nasal shedding compared to the WIV_VIC_ and unvaccinated groups over time (**Fig. 4A**). Area under the curve analysis (**Fig. 4B**) showed these differences to be significant and shedding in the WIV_Vic_ vaccinated pigs showed a similar high level to the naïve unvaccinated group. Samples obtained on 8 or 9 dpc at postmortem (PM), revealed that viral RNA could not be detected in nasal turbinate or the lower respiratory tract in any group (**Sup Fig. 1B**) indicating resolution of infection.

**FIG 4.**
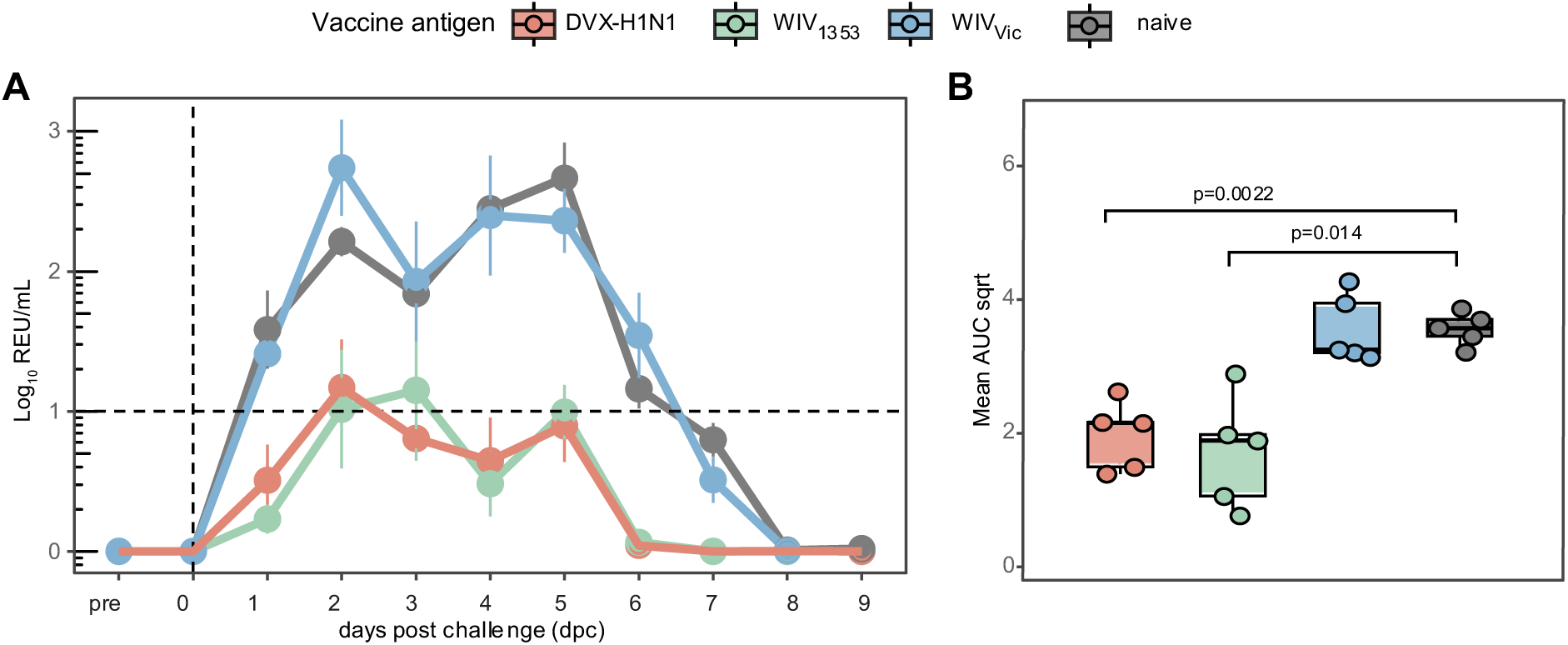
Nasal shedding of viral RNA in pigs and assessment of cell-mediated immune response. Viral RNA shedding following challenge with the swine-origin H1N1 virus A/swine/England/1353/2009 (1A.3.3.2) was assessed daily in nasal swabs by RRT-qPCR and is expressed as **(A)** mean log10 Relative Equivalent Units (REU)/ml (+/-SEM) over time (x-axis: days post challenge), and **(B)** square root of Area under the curve (AUC sqrt). For (B), p values were calculated using Wilcoxon signed-rank test. All p values are tabulated in **Supplementary Table 3.**

We delved deeper into identifying humoral immune responses in all pigs immunized with DVX-H1N1 by detecting influenza virus-specific serum antibody levels monitored longitudinally on 0, 28, 42, and 63dpi employing the HI assay, PV serum neutralising HA and NA inhibition assays, and an indirect ELISA to detect NP antibodies (**Fig. 5**).

**FIG 5.**
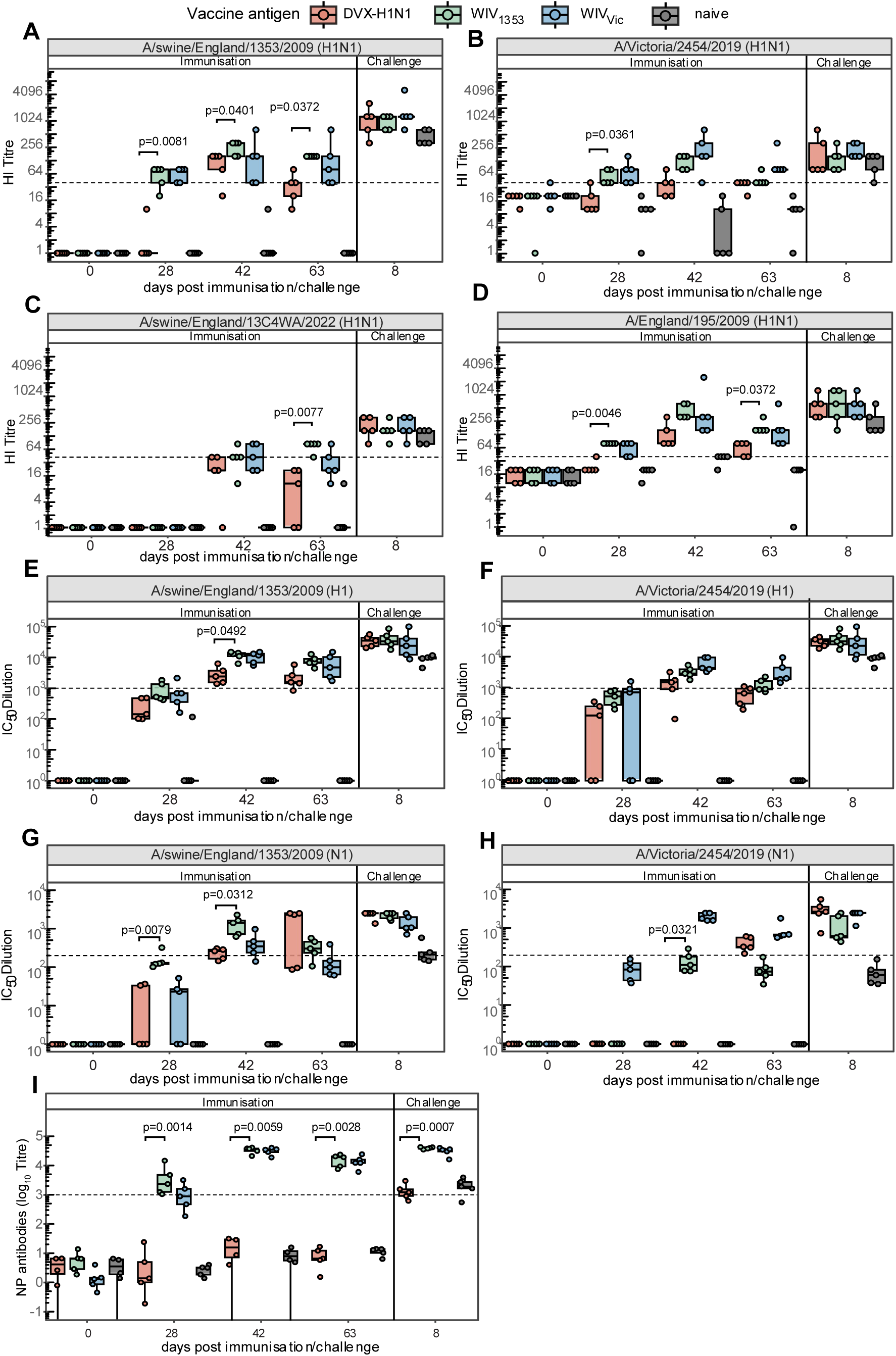
Pig humoral immune response monitoring for all groups of pigs. Serum samples obtained at 0 days post-immunization (dpi), 28dpi, 42dpi, 63dpi, and terminally at 8 days post-challenge (dpc) were analysed. Hemagglutination inhibition (HI) titres were measured using the H1N1 virus antigens (**A**) A/swine/England/1353/2009 (WIV_1353_), (**B**) A/Victoria/2454/2019 (WIV_Vic_), (**C**) a pig contemporary strain, A/swine/England/13C4WA/2022 and (**D**) A/England/195/2009. The dotted line indicates an HI titre of 40 which is considered seroprotective. Serum neutralising HA IC_50_ values using HA PV based on (**E**) A/swine/England/1353/2009 and (**F**) A/Victoria/2454/2019. NA inhibition IC_50_ values monitored respectively using NA PV based on (**G**) A/swine/England/1353/2009 and (**H**) A/Victoria/2454/2019. **(I)** Serum samples were assessed by an Influenza A virus nucleoprotein (NP) indirect ELISA (IDvet) for anti-NP antibody responses. A titre below 10^3^ is considered negative (dotted line). For all plots, the median and interquartile range of individual pig serum samples are shown (n=5 per vaccination group). *p*-values were determined via Kruskal allis test followed by post hoc Dunn’s test where *p<0.05* indicates a significant difference between groups. Only p values that are statistically significant between DVX-H1N1 and WIV_1353_ are indicated. All other p values are tabulated in **Supplementary Table 4**.

A panel of H1N1 2009 and contemporary human-and swine-origin 1A.3.3.2 strains representing 2009 and 2019 viruses were used in the HI assay (**Table 3**). Serological responses to DVX-H1N1 **(Fig. 5A-D)** showed HI titres increasing significantly from 28dpi to 42dpi against all virus antigens tested and the WIV_1353_ and WIV_VIC_ immunized groups had HI titres ≥40 by 28dpi, increasing modestly by 42dpi. As expected, the highest titres were elicited by the homologous antigens sw/EN/1353/09 and VIC/2454/19 in the WIV_1353_, and WIV_VIC_ groups respectively. Titres for all immunized groups were also higher against the 1A.3.3.2 EN/195/09 strain compared to the contemporary swine-origin strain sw/EN/13C4WA/2022. The humoral response in DVX-H1N1-immunized pigs reached the accepted seroprotective threshold (HI≥40) but overall had lower HI titres than the WIV vaccinated groups. Serum neutralizing HA titres monitored using PVs homologous to the control WIV vaccines showed a similar humoral response over time (**Fig. 5E,5F**) with the highest immunization phase titres observed at 42dpi in the immunized groups with minimum IC_50_ values of 1000. Similarly, NAI activity specific for A/swine/England/1353/2009 (N1 PV) and A/Victoria/2454/2019 (N1 PV) **(Fig. 5G,5H)** was detected. As expected, in all groups, titres were higher for homologous antigen-virus combinations compared to heterologous vaccination-virus combinations regardless of the viruses belonging to the same 1A.3.3.2 clade. Overall, the hemagglutination inhibition titres mirrored IC_50_ HA neutralising and NAI activity values. Additionally, during the immunization phase, elevated NP-specific antibody titres were detected in the WIV-vaccinated groups, but not in the group that received DVX-H1N1 which does not contain NP antigen (**Fig. 5I**) as expected. Such a distinction is advantageous for Differentiating Infected from VAccinated (DIVA) vaccine strategies (47).

Serological responses following challenge (8dpc) indicated that the HI antibody response to A/swine/England/1353/2009, the challenge strain, increased in all groups by 8dpc (**Fig. 5A-D**) suggesting successful seroconversion. The level of HI antibodies from all groups against all strains tested increased after virus inoculation, indicating an effective challenge (**Fig. 5A-D**). HA serum neutralising and NAI titres obtained using PV assays **(Fig. 5E-H)** showed similar patterns to the HI assays (**Fig. 5A-D**) with a marked increase in IC_50_ after challenge. For all groups, titres were higher for homologous antigen-virus combinations compared to heterologous (belonging to the same clade but drifted over time) vaccination-virus combinations. Notably, despite high anti-HA and anti-NA titres, the group immunized with the human whole inactivated vaccine (WIV_VIC_), representing a human seasonal vaccine stain from 2019 was unable to control the infection and reduce viral shedding (**Fig. 4A-B**) unlike the DVX-H1N1 vaccine, and similar to (no significant difference in viral shedding throughout the challenge window) with the naive negative control group. NP-reactive antibody levels remained high in the WIV-vaccinated groups and rose rapidly in the unvaccinated group; this was not observed in the DVX-H1N1 immunized pigs (**Fig. 5I**), because the infection was controlled in this group (**Fig. 4A**).

Cellular responses were evaluated by stimulating PBMCs from all immunized groups collected at different time points with the challenge strain WIV_1353_ or the WIV_Vic_ 2019 antigens. Stimulation by WIV_1353_ did not elicit any appreciable cellular response during the immunization window in all vaccine groups; however, at 8dpc, PBMC samples from three of the five DVX-H1N1 pigs produced >100 spot-forming colonies (SFC)/million cells compared to both WIV immunized groups (**Fig. 6A**). PBMC activation with WIV_VIC_ was apparent in the homologous WIV_VIC_ vaccine group during the immunization phase, and in WIV_1353_ pigs (**Fig. 6B**). At 8 dpc, PBMC samples from three of the five DVX-H1N1 immunized pigs reacted to the WIV_Vic_ 2019 antigen, suggesting that the challenge resulted in an increase in cellular responses not just against the challenge virus but against WIV_VIC_ as well (**Fig. 6**). The naive control group did not show increased spot-forming colonies (SFC) as expected, even after challenge.

**FIG 6.**
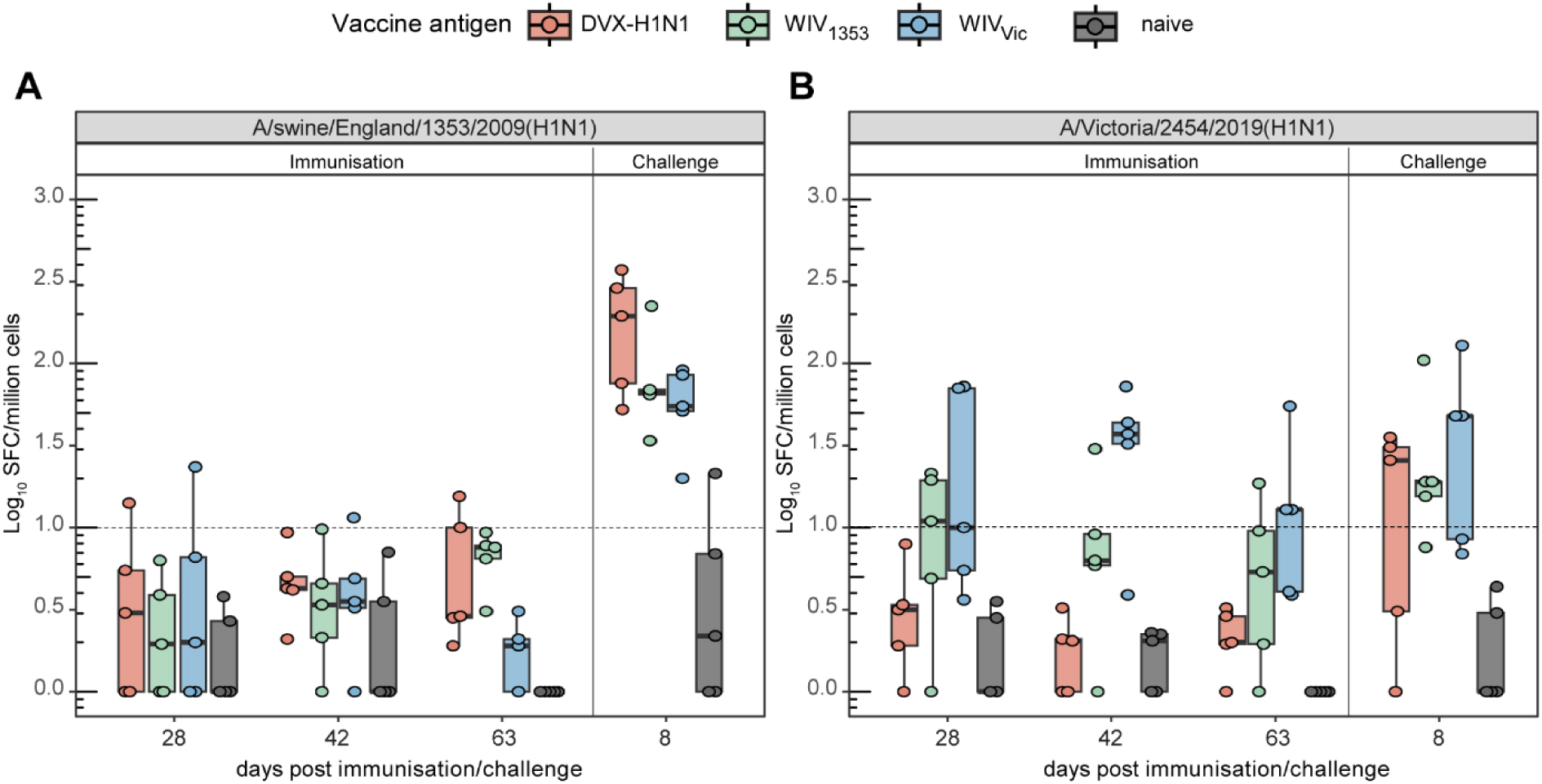
Pig cellular immune responses were also investigated by stimulating isolated PBMCs with inactivated whole inactivate virus (WIV) vaccine strains **(A)** WIV_1353_, and **(B)** WIV_Vic,_ to assess their IFN-γ response at various timepoints. Results are expressed as log_10_ transformed Spot Forming Cells (SFC)/million cells (+/-SEM). PMBC responses are shown for samples obtained after prime immunization (28dpi), 2 weeks after boost immunization (42dpi), 1 week prior to challenge (63dpi), and 1 week after challenge (8dpc). SFC below 1 Log_10_ is considered background. *p* values were determined via Kruskal allis test followed by post hoc Dunn’s test. There are no significant *p* values *(<0.05*) between DVX-H1N1 and WIV_1353_. All other p values are tabulated in **Supplementary Table 5**.

## DISCUSSION

Pandemic influenza remains one of the most serious threats to human and animal health worldwide. Swine influenza viruses contribute significantly to influenza virus reassortment through antigenic shift, leading to heightened zoonotic and pandemic risk (48). Here we presented new influenza vaccine technology by creating fully synthetic antigens that can elicit broad immune responses in different animal models as well as showing efficacy by reducing viral shedding in a pig model of influenza infection.

We assessed individual antigen immunogenicity in the mouse model and then the efficacy and immunogenicity of these antigens delivered as a single plasmid was evaluated in a well-established animal model of human influenza, the pig. Our findings demonstrate that digitally designed immune-optimized (DVX) antigens are immunogenic and provide highly effective immune protection against challenge with a relevant swine virus.

Mouse immunization with DVX-H1 resulted in H1 neutralizing titres IC_50_≥10^3^ (**Fig. 2A**) and hemagglutination inhibition titres ≥40 (**Fig. 2B**) that were higher than H1_ctrl_ against drifted H1 swine and human PV representatives and comparable against seasonal H1 PV. All neuraminidase antigens elicited poor levels of antibody-mediated inhibition against the N1 PV panel, except for DVX-N1 against EN/195/09 (N1) (**Fig. 2C**). Seroprotective HI titres elicited by DVX-H1 against two of the three swine A(H1)v viruses tested are promising and helps towards our goal of developing pandemic-proof vaccines. Given that HA contains the major immunodominant IAV epitopes and neutralization of HA is a major correlate of humoral immunity, these findings showing H1 cross-clade immune responses indicate the enhanced potential of DVX antigens to provide immune protection from strains undergoing antigenic drift and reassortment. This is a strong proof-of concept for *in-silico* designed antigens to offer broad protection against diverse variants that have the potential to emerge in future. Unlike existing strain-specific vaccines, our computationally engineered antigens induce broad neutralization within the H1 subtype encompassing human pandemic, seasonal, and zoonotic strains, thus mitigating the risk of a future swine influenza pandemic. As such, the data presented here (**Fig. 2**) also highlights the limitation of strain-specific antigens when it comes to pandemic preparedness.

Given that N1 and M2 are known to potentiate the immunogenicity of H1 in influenza vaccines (13,15,49–53), we therefore investigated the immunogenicity and efficacy of a DNA vaccine incorporating the computationally designed H1, N1, and M2 antigens (**Fig.1**) as described in mice (**Fig. 2**) in a more relevant animal model of influenza, the pig. The same broad HA neutralizing titres were also observed for candidates immunised with DVX-H1N1 in pigs. These titres, however, were higher in mice **(Fig. 2A)** compared to pigs **(Fig. 3B, 5B)** and can probably be attributed to several factors. Experimental mice are in-bred compared to the outdoor bred pigs (tested seronegative for influenza prior to the study) employed in the challenge; and both will have vastly different immune backgrounds. Antigens were also given individually in mice, as compared to three antigens in a single DNA-plasmid in pigs. We also opted for subcutaneous (SC) immunization in mice, and intradermal (ID) delivery via needle-free device in pigs. All these factors may have contributed to the limitations of our findings and are variables that can be further assessed in future iterations and improvements to these digitally designed antigens. For hemagglutination inhibition, the panel consisted of a pair of human isolates, A/Victoria/2454/2019 (H1N1), and a human seasonal/pandemic predecessor A/England/195/2009 (H1N1), a pair of swine isolates, A/swine/England/1353/2009 (H1N1 (vaccine), and contemporary strain A/swine/England/13C4WA/2022. Both WIV_1353_ and WIV_VIC_ showed higher HI titres than DVX-H1N1 (**Fig. 5A**), but these results were expected, as the DVX-H1N1 vaccine are not just HA head-specific but rather geared to be more broadly neutralizing towards Clade 1A HA antigens regardless of strains are human, swine, and/or contemporary. Additionally, improved neuraminidase inhibition by DVX-H1N1 was observed in pigs compared to DVX-N1 in mice (**Fig. 3C**, **5C**).

Overall, the immune responses observed in pigs during both the immunization and challenge windows were multi-faceted inducing both humoral, and cellular responses **(Fig. 3-6)**. Protective immunity correlated with HI, anti-HA neutralizing, and anti-NA inhibiting titres in pigs immunized with DVX-H1N1 and WIV_1353_ vaccines; however, this did not correlate for WIV_VIC_. Additionally, all these immune readouts markedly increased post-challenge for all vaccination groups indicating successful infection with A/swine/England/1353/2009 (H1N1) which boosted existing vaccine-induced immunity **(Fig. 3-6)**. This study demonstrates that immune correlates of protection extending beyond HI and neutralizing antibodies have conferred protection, reducing virus shedding; with novel vaccine antigens shown to recruit broader range of immunological effector mechanisms, important for pan-or universal influenza vaccine strategies.

Vaccines will continue to play an important role in containing pandemics. DVX technology as presented here has been developed to respond to high-priority virus threats with vaccines that can readily be deployed when the need arises, ensuring that the tools needed to protect health and respond to future pandemic threats are available.

## MATERIALS AND METHODS

### *In-silico* design of vaccine antigens

Nucleotide sequences of the hemagglutinin (HA) of pandemic lineage of the H1N1 subtype and neuraminidase (NA) of HxN1 subtype, were downloaded from the NCBI database. Utilizing the dataset of sequences of H1N1, and HxN1 collected from 1918 to 2018, unique antigen sequences were generated for HA, and NA, that were phylogenetically closest to all the input sequences in comparison to any sequence in the downloaded dataset. Briefly, the sequences were trimmed to the coding regions respectively for HA, and NA, and were filtered for redundancy at 95% nucleotide sequence identity. Multiple sequence alignment of HA, and NA, were generated using the MAFFT algorithm (54) with default parameters. Phylogenetic trees for HA, and NA, were produced using the previously generated multiple sequence alignment (MSA) as an input to the IQ-tree algorithm (55). The optimal nucleotide model for phylogenetic tree generation was chosen according to Bayesian information criteria (BIC) score. The phylogenetic tree and MSA were used as input to Hyphy (56) to generate antigens.

### Production of the challenge strain and whole inactivated virus (WIV) vaccines

Whole inactivated virus (WIV) vaccines were prepared from two H1N1 pandemic 2009 Clade 1A.3.3.2 viruses; A/swine/England/1353/2009 (57) strain and a representative human seasonal vaccine strain A/Victoria/2454/2019 (H1N1) (provided as the reassortant IVR-207), kindly provided by the WHO CC (The Francis Crick Institute, London, UK). Virus stocks were propagated in embryonated hens’ eggs and inactivated using beta-propiolactone at 1:2000 dilution for 2 hours as previously described (58). The resulting WIV_1353_ and WIV_VIC_ antigens were assessed using the Hemagglutination (HA) assay (59). Each 1.5 mL dose was standardized to contain 640 HA units (HAU) using endotoxin-free PBS (Gibco) and formulated as a 1:2 antigen:adjuvant mixture with TS6 adjuvant (Ceva) as described previously (44). The challenge strain, A/swine/England/1353/2009 (H1N1), was propagated in Madin-Darby canine kidney (MDCK_HK_) cells and titrated using the TCID_50_ method (59).

### Production of DNA vaccines and lentiviral pseudotype viruses (PV)

Selected antigen sequences were codon-optimized for human expression via the GeneOptimizer algorithm (60) and were then synthesized and cloned into pEVAC (GeneArt, Germany). Cloned DNA was used for the production of DNA vaccines and controls (**Table 1**).

H1 and N1 sequences (**Table 2**) were produced as HA or NA lentiviral pseudotype viruses (PV) as described previously (61,62) via FuGENE^®^ HD (ProMega, USA) transfection of HEK 293T/17 cells in 6-well plates with four plasmids: pEVAC encoding HA for HA PV, and HA and NA for NA PV, pCSFLW encoding a luciferase reporter gene, p8.91 gag-pol encoding lentiviral G antigen and polymerase and a type II transmembrane protease serine 4 (TMPRSS4). Twenty-four hours post-transfection for HA pseudotype viruses, 1 unit of exogenous neuraminidase in 50 µL PBS (Sigma Aldrich N2876, Paisley, UK) was added per well. Supernatant was collected 48 hours post-transfection, passed through a 0.45 µm filter, and stored at-80°C. Titration was carried out as described previously (61,62).

### Ethics

*In vivo* studies were conducted in accordance with UK Home Office regulations under the Animals (Scientific Procedures) Act 1986 (ASPA) following ethical approval. Mouse studies were approved by the Animal Welfare Ethical Review Body (AWERB), University of Cambridge (Project license P8143424B). Pig study number PP9878849-2-002 was approved by the AWERB of the Animal and Plant Health Agency (APHA). Results are reported according to the ARRIVE Guidelines (63).

### Vaccine Immunogenicity in the mouse model

Six to eight-week-old female BALB/c mice were obtained from Charles River Laboratories and housed in 7 groups of 6 mice each at the University Biomedical Services, University of Cambridge. DNA at a concentration of 50 µg DNA suspended in 50 µL of endotoxin-free PBS (Gibco) was administered subcutaneously (SC) on the rear flank of each mouse. Following the initial prime at 0 days post-immunization (dpi), immunizations were boosted on 14, 28, and 42dpi. The respective groups (**Table 1**) received DNA encoding a DVX antigen HA (H1), NA (N1), or M2 (M2) or the respective control immunogens HA (H1_ctrl_), NA (N1_ctrl_), or M2 (M2_ctrl_) derived from specific viruses. One group was injected with 50 µL of plasmid pEVAC (no insert) as negative control. Blood samples were collected on 42 and 56dpi. All mice were euthanized under non-recovery isofluorane (5%) anesthesia on 70dpi and terminal blood samples were collected. Serum samples were kept at-20°C for later determination of the humoral immune responses induced by vaccination.

### Immunogenicity and challenge study in pigs

Five-week-old Landrace cross male pigs (n=20) were obtained from a commercial high health status herd and housed in 4 groups (n=5 per group). Pigs were screened for absence of influenza A viral RNA by matrix gene RRT-qPCR (64) and absence of influenza A antibodies to H1N1 1A.3.3.2 antigens by Hemagglutination Inhibition (HI) (65,66). Throughout the study all pigs received weaner/grower feed and had access to water ad-libitum. Pigs were vaccinated (0 dpi) at 7 weeks of age and boosted 28 days later. The DVX-H1N1 DNA vaccine was administered intradermally (ID) using a PharmaJet® Tropis device (Pharmajet) by delivery of four doses (two doses posterior to both left and right ears). The monovalent WIV_1353_ or WIV_Vic_ vaccines were administered intramuscularly (IM) with two doses delivered into the trapezius muscle 25-30mm posterior to each ear. A negative naïve control group was not vaccinated (**Table 1**).

Pigs were challenged intranasally (IN) on 69dpi with 4 mL (2 mL per nostril) of A/swine/England/1353/2009 (H1N1) virus delivered as an atomized spray of droplets with diameters ranging from 30-100 µm using a MAD300 device (Teleflex, USA). Back-titration of the challenge stock confirmed that 1.67×10^6^ TCID_50_ was administered to each pig. Animals were observed daily post-challenge (dpc) for signs of illness and/or welfare impairment using a clinical scoring system which includes demeanor, appetite, rectal temperature and respiratory signs such as coughing and sneezing (67). Clotted and heparin anticoagulated blood samples (BD BioSciences SST and Heparin vacutainers) were obtained prior to immunization, at 28, 42, and 63dpi as well as after challenge at 8 days post challenge (dpc). Serum samples obtained from clotted blood were archived at-20°C and PBMCs were isolated from anticoagulated blood and cryopreserved as described previously (44). Nasal swabs (one per nostril) were obtained from each pig prior to vaccination and challenge and then daily post-challenge until the end of the study and stored dry at-80°C (44). The pigs were humanely euthanized with an overdose of intravenous pentobarbital sodium at the study end on 8dpc for WIV-vaccinated groups, and 9dpc for DVX-H1N1 and negative control groups. Bronchioalveolar lavage (BAL) samples were obtained *postmortem* in Dulbecco’s PBS (Gibco) and cellular and supernatant fractions were archived as described above.

### Real-time reverse transcription polymerase chain reaction (RT-qPCR)

To evaluate viral RNA shedding by RRT-qPCR both nasal swabs obtained at specified timepoints from each pig were immersed in 1 mL of Leibovitz L-15 medium (ThermoFisher Scientific), containing 1% fetal bovine serum (FBS) and 1% penicillin/streptomycin (Gibco, 5000 U/mL). Total RNA was extracted from the swab suspensions and bronchoalveolar lavage fluid (BAL) using the RNeasy® Mini kit (Qiagen, Crawley, UK) according to the manufacturer’s instructions. Viral RNA was quantified by RRT-qPCR directed against the swine IAV M-gene (44,68). The amount of RNA present is expressed as relative equivalent units (REU) of RNA using a standard 10-fold dilution series of RNA purified from the same batch of virus used for challenge, of known TCID_50_ titre. Although these units measure the amount of viral RNA present and not infectivity, it may be inferred from the linear relationship with the dilution series that they are proportional to the amount of infectious virus present, as described previously (44).

### Humoral immune response

Serum neutralizing titres against a H1 PV panel (**Table 2**) were determined for both mice and pigs via pseudotype microneutralization assay (pMN) (61). For mice, serum samples were diluted 1:50, and for pigs 1:100, responses were normalized against virus only (0% neutralization) and cell only (100% neutralization) controls and plotted on GraphPad Prism 10.3.1. Values were fitted using nonlinear regression (curve fit) where x is log fold-dilution of sera and y is normalized response. Results are calculated as IC_50_ fold dilution of sera that resulted in 50% neutralization of virus.

Neuraminidase inhibition titres against an N1 PV panel (**Table 2**) were determined for both mice and pigs via pseudotype Enzyme Linked Lectin Assay (pELLA) (62). Mouse and pig sera were initially diluted 1:20 and then serially diluted two-fold in duplicate. The IC_50_ was calculated as the inverse dilution of serum or antibody concentration that resulted in 50% inhibition of NA activity as determined via GraphPad Prism 10.3.1.

Mouse and pig antisera were assessed by HI using an H1N1 reference virus panel (**Table 3**). HI titres of 40 and above are considered positive for mouse and pig antisera. Additionally, pig antibody levels were measured longitudinally using an indirect multi-species ELISA to detect the nucleoprotein (NP) of influenza A virus (ID Screen® IDvet).

### Cellular immune Response (Porcine IFN-γ ELISpot assay)

PBMC immune stimulation resulting from immunization and/or challenge was assessed using an ELISpot assay (69) stimulated with WIV_1353_ and WIV_Vic_ antigens used as vaccines. The number of Spot Forming Cells (SFC) per 10^6^ cells was counted with an automated ELISpot reader (AID) and corrected for background (tissue culture medium-stimulated). Results are given for samples in which cells reacted to mitogen (Pokeweed, Sigma) as a positive control for the functional capability of cells to produce IFN-γ.

## Statistical analysis

All statistical analyses were performed with RStudio for Windows (RStudio 2024.09.0 Posit Software, PBC). For both animal studies, a Wilcoxon signed-rank test or a Kruskal Wallis test followed by post hoc Dunn’s test were used to determine if there were statistically significant differences between two or more groups in comparison to a control group. Only statistically significant p-values are displayed.

## DATA AVAILABILITY

All data supporting the findings of this study are available within the paper and its Supplementary Information.

## ACKNOWLEDGMENTS

We gratefully acknowledge the excellent animal care provided for pigs by the APHA Animal Sciences Department. We are also grateful to the Worldwide Influenza Centre (WIC), The Francis Crick Institute for provision of the reassortant strain A/Victoria/2454/2019 (IVR-207).

## AUTHOR CONTRIBUTIONS

J.M.D., S.B.S., and S.K.A. produced DNA for immunization and pseudotype viruses, performed *in vitro* assays, pMN, pELLA, and Luminex binding. G.W.C., P.T., and J.M.D. performed the mouse immunogenicity studies. J.M. performed HAI in mice. S.F. designed the vaccine antigens.

B.A. performed purification and preparation of DNA for antigen selection. N.T. and J.M. provided key reagents. J.M.D., P.V.D., and M.D. analysed the data. S.V., S.K.A., B.S., and M.D. produced all figures. S.K.A. and M.D. performed statistical analyses. R.K., R.W., and J.L.H. secured the funding. J.L.H conceptualised the investigation. J.M.D., G.W.C, P.V.D, H.E., and J.L.H designed the experiments. H.E. and J.L.H. supervised the project. J.M.D., P.V.D., and S.B.S. wrote the original draft. G.W.C, J.M., R.W., H.E., and J.L.H reviewed all data and edited the manuscript. All authors reviewed and approved the manuscript.

## COMPETING INTERESTS

J.M.D., S.B.S., S.K.A., B.S., G.W.C., M.D., R.K., R.W., and J.L.H. are employees or shareholders of DIOSynVax Ltd. S.F. is an employee of Microsoft. The sequences of the DVX antigens have been patented under UK Patent Application No. GB2414517.8, Influenza vaccines, PCT/GB2022/052534, and PCT/GB2024/052670 (Influenza Antigen Synergies patent).

## FUNDING

This research was funded by Bill and Melinda Gates Foundation: Grand Challenges Universal Influenza Vaccines Award: Ref: G101404 to J.L.H and Innovate UK, UK Research and Innovation (UKRI), for the project: Digital Immune Optimized and Selected Pan-Influenza Vaccine Antigens (DIOS-PIVa) Award Ref: 105078 to J.L.H. Influenza research at APHA is supported by Defra and the devolved Scottish and Welsh Governments previously through FluFutures2 (SE2213) and currently through FluFocus (SE2227).

