## Supplementary file for "Immunogenicity and Efficacy of Digitally Immune Optimised H1N1 Vaccine Candidates in Swine and Murine Animal Models"

### SUPPLEMENTARY MATERIAL

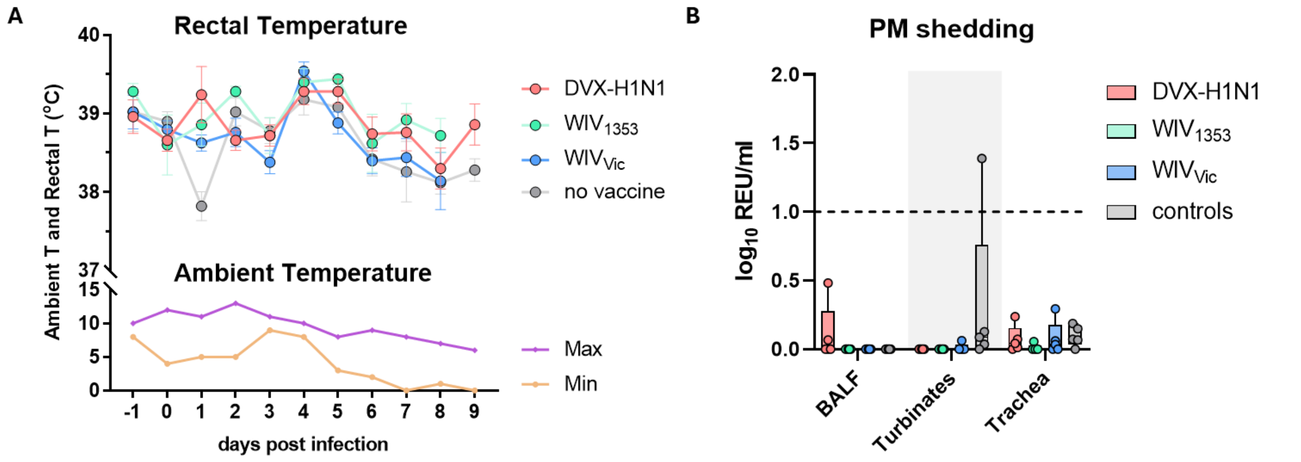


**Supplementary Figure 1. (A)** Pig temperature monitoring during challenge window. Daily rectal and ambient temperatures (°C) were recorded prior to and on each day post-challenge as indicated. **(B)** Quantification of viral RNA in bronchoaveolar lung lavage fluid (BALF) and swabs of respiratory tract tissues (nasal turbinate or trachea) obtained following Postmortem (PM) at 8 or 9 days post-challenge.

**Supplementary Table 1.** *p* values as calculated using Wilcoxon signed-rank test between groups of mice immunized with: (**A,B**) DVX-H1, $H1_{ctrl}$, and pEVAC for **Fig. 2A-B**; and (**C**) DVX-N1, $N1_{ctrl}$, and pEVAC for **Fig. 2C**. *p* values that cannot be computed (dataset values=0) are designated as >1.

| **H1 PV (Fig. 2A)** | DVX-H1 vs $H1_{ctrl}$ | DVX-H1 vs pEVAC | $H1_{ctrl}$ vs pEVAC |
|---|---|---|---|
| SC/1/1918 | 0.0037 | 0.0055 | 0.4652 |
| CA/07/09 | 0.3611 | 0.0635 | 0.0587 |
| BR/59/07 | 0.0055 | 0.0028 | >1 |
| MI/45/15 | 0.0591 | 0.0256 | 0.0469 |
| BR/2/18 | 0.2539 | 0.0256 | 0.0231 |
| VIC/2454/19 | 0.1198 | 0.0027 | 0.0039 |
| VIC/2570/19 | 0.4286 | 0.0028 | 0.0039 |
| WAS/19/20 | 0.6664 | 0.0038 | 0.0038 |
| sw/EN/1353/09 | 0.0039 | 0.0013 | 0.0150 |
| sw/GX/13 | 0.0013 | 0.0013 | >1 |
| sw/HN/18 | 0.0028 | 0.0256 | >1 |
| **H1N1 Virus antigen (Fig. 2B)** | DVX-H1 vs $H1_{ctrl}$ | | |
| CA/07/09 | 0.6842 | | |
| MI/45/15 | 0.0836 | | |
| BR/2/18 | 0.0585 | | |
| HUN/42443/15 | 0.0740 | | |
| HEB-HAI/1572/19 | 0.0740 | | |
| HESS/47/20 | >1 | | |
| **N1 PV (Fig. 2C)** | DVX-N1 vs $N1_{ctrl}$ | DVX-N1 vs pEVAC | $N1_{ctrl}$ vs pEVAC |
| EN/195/09 | 0.0028 | 0.0028 | >1 |
| BR/2/18 | 0.0284 | >1 | 0.0284 |
| VIC/2454/19 | 0.0898 | 0.4047 | 0.0284 |
| sw/EN/1353/09 | 0.4047 | >1 | 0.4047 |
| sw/NC/20 | >1 | >1 | >1 |

**Supplementary Table 2.** *p* values as calculated using Wilcoxon signed-rank test between groups of pigs immunized with DVX-H1N1, $WIV_{1353}$, $WIV_{VIC}$, and naive for **Fig. 3**. *p* values that cannot be computed (dataset values=0) are designated as >1.

| **H1 PV (Fig. 3B)** | DVX-H1N1 vs $WIV_{1353}$ | DVX-H1N1 vs $WIV_{VIC}$ | DVX-H1N1 vs naive | $WIV_{1353}$ vs $WIV_{VIC}$ | $WIV_{1353}$ vs naive | $WIV_{VIC}$ vs naive |
|---|---|---|---|---|---|---|
| SC/1/1918 | 0.0670 | 0.4206 | 0.0075 | 0.0112 | 0.0067 | 0.0075 |
| CA/07/09 | 0.5258 | 0.6004 | 0.0075 | 0.5258 | 0.0073 | 0.0075 |
| EN/195/09 | 0.0112 | 0.3095 | 0.0075 | 0.0670 | 0.0067 | 0.0075 |
| BR/59/07 | 0.7972 | >1 | 0.1797 | 0.7972 | 0.4237 | 0.1797 |
| MI/45/15 | 0.0556 | 0.4206 | 0.0075 | 0.3457 | 0.0075 | 0.0075 |
| BR/2/18 | >1 | 0.4237 | 0.0040 | 0.4237 | 0.0040 | 0.0056 |
| VIC/2454/19 | 0.0079 | 0.0112 | 0.0075 | 0.0907 | 0.0075 | 0.0067 |
| VIC/2570/19 | 0.0556 | 0.0119 | 0.0075 | 0.0937 | 0.0075 | 0.0073 |
| WAS/19/20 | 0.0311 | 0.3465 | 0.0056 | 0.2087 | 0.0075 | 0.0073 |
| sw/EN/1353/09 | 0.0200 | 0.0212 | 0.0075 | 0.6558 | 0.0067 | 0.0073 |
| sw/GX/13 | 0.4237 | >1 | >1 | 0.4237 | 0.4237 | >1 |
| **N1 PV (Fig. 3C)** | | | | | | |
| EN/195/09 | 0.0159 | 0.1508 | 0.0075 | 0.0317 | 0.0075 | 0.0075 |
| BR/2/18 | 0.0079 | 0.0075 | 0.0075 | 0.0075 | 0.0075 | 0.0040 |
| VIC/2454/19 | 0.0079 | 0.0119 | 0.0254 | 0.0119 | 0.0075 | 0.0073 |
| sw/EN/1353/09 | 0.0079 | 0.4206 | 0.0075 | 0.0317 | 0.0075 | 0.0075 |
| sw/NC/20 | 0.2222 | 0.0345 | 0.0075 | 0.1388 | 0.0075 | 0.1797 |

**Supplementary Table 3.** *p* values as calculated by post hoc Dunn's test between groups of pigs immunized with DVX-H1N1, $WIV_{1353}$, $WIV_{VIC}$, and naive for **Fig. 4**. Only comparisons that were found significant via Kruskal Wallis analysis were subjected to post hoc tests; non-significant comparisons are labelled ns. *p* values that cannot be computed (dataset values=0) are designated as >1. *p* values for **Fig. 4B** were calculated using Wilcoxon signed-rank test.

| **Fig. 4A** | | | | | | |
|---|---|---|---|---|---|---|
| **Days post challenge** | DVX-H1N1 vs $WIV_{1353}$ | DVX-H1N1 vs $WIV_{VIC}$ | DVX-H1N1 vs naive | $WIV_{1353}$ vs $WIV_{VIC}$ | $WIV_{1353}$ vs naive | $WIV_{VIC}$ vs naive |
| 1 | 0.4227 | 0.0777 | 0.0325 | 0.0103 | 0.0033 | 0.7083 |
| 2 | 0.7083 | 0.0046 | 0.0422 | 0.0139 | 0.0975 | 0.4227 |
| 3 | ns | ns | ns | ns | ns | ns |
| 4 | 0.7483 | 0.0284 | 0.0120 | 0.0120 | 0.0046 | 0.7483 |
| 5 | 0.9574 | 0.0139 | 0.0033 | 0.0162 | 0.0039 | 0.6305 |
| 6 | 0.7848 | 0.0022 | 0.0074 | 0.0053 | 0.0163 | 0.7022 |
| 7 | >1 | 0.0304 | 0.0019 | 0.0304 | 0.0019 | 0.3492 |
| **Fig. 4B** | | | | | | |
| | 0.5476 | 0.0079 | 0.0022 | 0.0079 | 0.014 | 0.8413 |

**Supplementary Table 4.** *p* values as calculated by post hoc Dunn's test between groups of pigs immunized with DVX-H1N1, $WIV_{1353}$, $WIV_{VIC}$, and naive for **Fig. 5**. Only comparisons that were found significant via Kruskal Wallis analysis were subjected to post hoc tests; non-significant comparisons are labelled ns. *p* values that cannot be computed (dataset values=0) are designated as >1.

| **Fig. 5A** A/swine/England/1353/2009 (H1N1) | | | | | | |
|---|---|---|---|---|---|---|
| **Timepoint** | DVX-H1N1 vs $WIV_{1353}$ | DVX-H1N1 vs $WIV_{VIC}$ | DVX-H1N1 vs naive | $WIV_{1353}$ vs $WIV_{VIC}$ | $WIV_{1353}$ vs naive | $WIV_{VIC}$ vs naive |
| 28dpi | 0.0081 | 0.0057 | 0.7758 | 0.9093 | 0.0033 | 0.0023 |
| 42dpi | 0.1532 | 0.7006 | 0.0321 | 0.2966 | 0.0004 | 0.0115 |
| 63dpi | 0.0410 | 0.1908 | 0.1078 | 0.4618 | 0.0003 | 0.0035 |
| 8dpc | ns | ns | ns | ns | ns | ns |
| **Fig. 5B** A/Victoria/2454/2019 (H1N1) | | | | | | |
| 28dpi | 0.0361 | 0.0121 | 0.5259 | 0.6791 | 0.0063 | 0.0017 |
| 42dpi | 0.1040 | 0.0395 | 0.1588 | 0.6646 | 0.0024 | 0.0005 |
| 63dpi | 0.5212 | 0.0239 | 0.0742 | 0.1057 | 0.0152 | 0.0001 |
| 8dpc | ns | ns | ns | ns | ns | ns |
| **Fig. 5C** A/swine/England/13C4WA/2022 (H1N1) | | | | | | |
| 28dpi | ns | ns | ns | ns | ns | ns |
| 42dpi | 0.3202 | 0.2694 | 0.0604 | 0.9121 | 0.0041 | 0.0029 |
| 63dpi | 0.0077 | 0.1614 | 0.3796 | 0.2066 | 0.0004 | 0.0227 |
| 8dpc | ns | ns | ns | ns | ns | ns |
| **Fig. 5D** A/England/195/2009 (H1N1) | | | | | | |
| 28dpi | 0.0046 | 0.0274 | 0.5667 | 0.5286 | 0.0007 | 0.0055 |
| 42dpi | 0.0561 | 0.1997 | 0.0960 | 0.5303 | 0.0004 | 0.0032 |
| 63dpi | 0.0372 | 0.1247 | 0.1247 | 0.5835 | 0.0003 | 0.0021 |
| 8dpc | ns | ns | ns | ns | ns | ns |
| **Fig. 5E** A/swine/England/1353/2009 (H1) | | | | | | |
| 28dpi | 0.1197 | 0.1630 | 0.1197 | 0.8721 | 0.0019 | 0.0032 |
| 42dpi | 0.0492 | 0.0799 | 0.1457 | 0.8294 | 0.0006 | 0.0013 |
| 63dpi | 0.0755 | 0.1961 | 0.0950 | 0.6279 | 0.0006 | 0.0031 |
| 8dpc | 0.9149 | 0.5212 | 0.0103 | 0.4543 | 0.0075 | 0.0543 |
| **Fig. 5F** A/Victoria/2454/2019 (H1) | | | | | | |
| 28dpi | 0.1166 | 0.2870 | 0.2393 | 0.6140 | 0.0060 | 0.0250 |
| 42dpi | 0.2153 | 0.0205 | 0.1314 | 0.2812 | 0.0060 | 0.0001 |
| 63dpi | 0.3599 | 0.0312 | 0.0950 | 0.2154 | 0.0097 | 0.0001 |
| 8dpc | 0.8307 | 0.5565 | 0.0120 | 0.4227 | 0.0064 | 0.0543 |

| Timepoint | DVX-H1N1 vs $WIV_{1353}$ | DVX-H1N1 vs $WIV_{VIC}$ | DVX-H1N1 vs naive | $WIV_{1353}$ vs $WIV_{VIC}$ | $WIV_{1353}$ vs naive | $WIV_{VIC}$ vs naive |
|---|---|---|---|---|---|---|
| **Fig. 5G** A/swine/England/1353/2009 (N1) | | | | | | |
| 28dpi | 0.0079 | 0.7534 | 0.3608 | 0.0192 | 0.0004 | 0.2195 |
| 42dpi | 0.0312 | 0.5536 | 0.0755 | 0.1183 | 0.0001 | 0.0178 |
| 63dpi | 0.8716 | 0.1960 | 0.0015 | 0.2579 | 0.0026 | 0.0593 |
| 8dpc | 0.8037 | 0.1591 | 0.0009 | 0.2462 | 0.0022 | 0.0567 |
| **Fig. 5H** A/Victoria/2454/2019 (N1) | | | | | | |
| 28dpi | >1 | 0.0004 | >1 | 0.0004 | >1 | 0.0004 |
| 42dpi | 0.0321 | 0.0004 | >1 | 0.1532 | 0.0321 | 0.0004 |
| 63dpi | 0.1614 | 0.2154 | 0.0060 | 0.0083 | 0.1782 | 0.0001 |
| 8dpc | 0.0950 | 0.6666 | 0.0007 | 0.2154 | 0.0848 | 0.0031 |
| **Fig. 5I** | | | | | | |
| 28dpi | 0.0036 | 0.0059 | 0.7483 | 0.8726 | 0.0095 | 0.0150 |
| 42dpi | 0.0102 | 0.0088 | 0.8725 | 0.9573 | 0.0064 | 0.0054 |
| 63dpi | 0.0074 | 0.0032 | 0.7890 | 0.7890 | 0.0160 | 0.0074 |
| 8dpc | 0.0006 | 0.0028 | 0.1088 | 0.6689 | 0.0692 | 0.1646 |

**Supplementary Table 5.** *p* values as calculated by post hoc Dunn's test between groups of pigs immunized with DVX-H1N1, $WIV_{1353}$, $WIV_{VIC}$, and naive for **Fig. 6** Only comparisons that were found significant via Kruskal Wallis analysis were subjected to post hoc tests; non-significant comparisons are labelled ns. *p* values that cannot be computed (dataset values=0) are designated as >1.

| **Fig. 6A** A/swine/England/1353/2009 (H1N1) | | | | | | |
|---|---|---|---|---|---|---|
| **Days post challenge** | DVX-H1N1 vs $WIV_{1353}$ | DVX-H1N1 vs $WIV_{VIC}$ | DVX-H1N1 vs naive | $WIV_{1353}$ vs $WIV_{VIC}$ | $WIV_{1353}$ vs naive | $WIV_{VIC}$ vs naive |
| 28dpi | ns | ns | ns | ns | ns | ns |
| 42dpi | ns | ns | ns | ns | ns | ns |
| 63dpi | 0.6228 | 0.0955 | 0.0058 | 0.0309 | 0.0011 | 0.2743 |
| 8dpc | 0.3358 | 0.1813 | 0.0008 | 0.7082 | 0.0161 | 0.0422 |
| **Fig. 6B** A/Victoria/2454/2019 (H1N1) | | | | | | |
| 28dpi | 0.1614 | 0.0407 | 0.4508 | 0.5181 | 0.0312 | 0.0051 |
| 42dpi | 0.1120 | 0.0033 | >1 | 0.1780 | 0.1120 | 0.0033 |
| 63dpi | 0.3528 | 0.0460 | 0.1634 | 0.2865 | 0.0202 | 0.0007 |
| 8dpc | 0.5729 | 0.2951 | 0.0679 | 0.6289 | 0.0169 | 0.0041 |
